# Microbial Community Dynamics Provide Evidence for Hypoxia During a Coral Reef Mortality Event

**DOI:** 10.1101/2022.02.24.481904

**Authors:** Shawn M. Doyle, Miabel J. Self, Joseph Hayes, Kathryn E.F. Shamberger, Adrienne M.S. Correa, Sarah W. Davies, Lory Z. Santiago-Vázquez, Jason B. Sylvan

## Abstract

In July 2016, a severe coral reef invertebrate mortality event occurred approximately 200km southeast of Galveston, Texas at the East Flower Garden Bank wherein upwards of 80% of corals in a 0.06 km^2^ area died. Based on surveys of dead corals and other invertebrates shortly after this mortality event, responders hypothesized that localized hypoxia was the most likely direct cause. However, no dissolved oxygen data were available to test this hypothesis because oxygen is not continuously monitored within the Flower Garden Banks sanctuary. Here we quantify microbial plankton community diversity based on four cruises over two years at the Flower Garden Banks, including a cruise just 5-8 days after the mortality event was first observed. In contrast with observations collected during baseline conditions, microbial plankton communities in the thermocline were differentially enriched with taxa known to be active and abundant in oxygen minimum zones or that have known adaptations to oxygen limitation shortly after the mortality event (e.g. SAR324, *Thioglobaceae, Nitrosopelagicus*, and *Thermoplasmata* MGII). Unexpectedly, these enrichments were not localized to the East Bank, but were instead prevalent across the entire study area, suggesting there was a widespread depletion of dissolved oxygen concentrations in the thermocline around the time of the mortality event. Hydrographic analysis revealed the southern East Bank coral reef (where the localized mortality event occurred) was uniquely within the thermocline at this time. Our results demonstrate how temporal monitoring of microbial communities can be a useful tool to address questions related to past environmental events.

**IMPORTANCE:** In the northwestern Gulf of Mexico in July 2016, upwards of 80% of corals in a small area of the East Flower Garden Bank coral reef suddenly died without warning. Oxygen depletion is believed to have been the cause. However, there was considerable uncertainty as no oxygen data is available from the time of the event. Microbes are sensitive to changes in oxygen and can be used as bioindicators of oxygen loss. In this study, we analyze microbial communities in water samples collected over several years at the Flower Garden Banks, including shortly after the mortality event. Our findings indicate that compared to normal conditions, oxygen depletion was widespread in the deep-water layer during the mortality event. Hydrographic analysis of water masses further revealed some of this low oxygen water likely upwelled onto the coral reef.

## 1. INTRODUCTION

In late July 2016, a severe mortality event occurred in a localized area of the East Bank coral reef within the Flower Garden Banks National Marine Sanctuary. During the mortality event, approximately 82% of coral colonies in a 0.06 km^2^ area of the East Bank (2.6% of the coral reef) experienced partial or full mortality (1). Many other sessile or sedentary invertebrates were affected as well including sponges, echinoderms, crustaceans, and mollusks. Within the affected area, mortality was concentrated in sand flats, channels, and other reef depressions and sometimes formed visible “bathtub rings”, with tissue above the rings appearing healthy, whereas tissue below exhibited bleaching, sloughing, and death (Fig. 1)(1).

**Figure 1.**
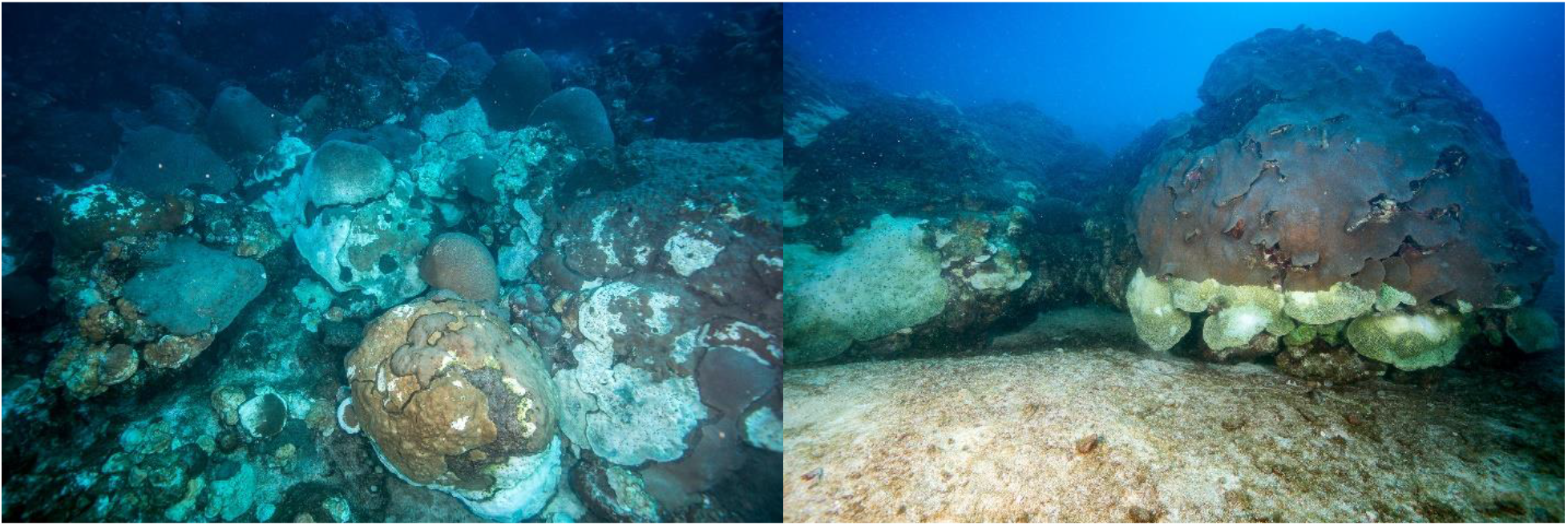
Corals affected by the 2016 localized mortality event at the East Flower Garden Bank. Photo Credit: G.P. Schmahl, NOAA-FGBNMS.

The localized mortality event is hypothesized to have been caused by the formation of hypoxia (defined as a dissolved oxygen (DO) concentration less than 2 mg/L) within reef depressions (1), suffocating animals in these areas. Unfortunately, no DO data are available from within the sanctuary at the time the mortality event occurred. As such, the presence and distribution of hypoxia within the Flower Garden Banks during the mortality event remains hypothetical. Despite this uncertainty, multiple mechanisms for how hypoxia may have formed on the East Bank coral reef have been proposed.

One proposed mechanism is an unusually large plume of coastal floodwater that transited over the Flower Garden Banks in the summer of 2016 provided favorable conditions for the local formation of hypoxic waters on top of the East Bank coral reef (2). Oceanographic and satellite data from the summer of 2016 showed the presence of turbid, brackish floodwater in the northwest Gulf of Mexico that was associated with bottom water hypoxia near the Texas-Louisiana coast (>100km from the Flower Garden Banks) in June 2016 (2). This plume of coastal water primarily originated from the Mississippi/Atchafalaya River system (3) and was supplemented by an unusually high (~10 times larger than usual) discharge of water from the Trinity, San Jacinto, and Brazos Rivers resulting from intense precipitation and flooding events (e.g. the “Tax Day Flood”) that occurred across southeast Texas and southwest Louisiana in the spring of 2016 (4). Upwelling-favorable (southerly) winds along the Texas coast in June and July 2016 subsequently drove this brackish floodwater plume offshore and over the coral reefs within the Flower Garden Banks (2). Microbial respiration of organic matter deposited by the plume has been hypothesized to have depleted dissolved oxygen concentrations within reef depressions, leading to invertebrate mortality (3).

Another proposed mechanism is that, in addition to the presence of a surface floodwater plume, dense water may have upwelled from the seaward side of the East Bank onto the coral reef and settled in reef depressions (3). Indeed, observations of water as far as 200m deep upwelling onto the East Bank have been observed before (5). This hypothesis is based on the principal that the formation of bottom water hypoxia is a vertical process which intrinsically depends not only on increased net respiration but also the formation of a vertically stratified water column (6, 7). This stratification necessarily inhibits exchanges between the water column and therefore prevents replenishment of consumed oxygen in the bottom boundary layer, in this case, depressions on the East Bank reef (8, 9). Higher water density, dissolved inorganic carbon (DIC), ammonium concentrations, and salinity, along with lower temperature and aragonite saturation state (Ω_ar_) were measured at 75 m near East Bank in August 2016, all of which support upwelling of water from the deeper Gulf of Mexico (GoM) onto the East Bank coral reef. Isopycnal direction and slope were also indicative of upwelling (3).

Microbial communities are key drivers of biogeochemical cycles in the oceans and respond quickly and specifically to environmental disturbances. As a result, shifts in microbial composition and structure can serve as useful bioindicators of specific biogeochemical changes (10), including the development and presence of hypoxia (11). Following the 2016 mortality event at East Bank, analysis of microbial communities in the overlying water column revealed community differences in the floodwater plume over East Bank compared to surface water outside the plume. In addition, deep-water communities were most abundant in subsurface waters at stations near East Bank where the localized mortality event occurred (3). Although these microbial data were interpreted as potential additional evidence of upwelling onto the East Bank, it was unknown if this finding was anomalous compared to normal conditions.

Here, we present new data from an additional three cruises that provide a baseline for the region in summer and fall. We test whether microbial communities collected shortly after the mortality event differed in composition and structure as compared with those collected during normal conditions in the same season within the Flower Garden Banks. We hypothesize that the presence of hypoxic waters on the Flower Garden Banks would be reflected in microbial plankton community composition and structure and allow us to further resolve what caused the mortality event. By establishing baseline microbial trends over time, we found the mortality event indeed coincided with significant differences in microbial community composition and structure. These differences were primarily due to the relative outgrowth of several microbial taxa typically found in marine oxygen minimum zones (OMZ), providing the first direct evidence of low-oxygen conditions around the time of the localized mortality event compared to baseline conditions. This microbial evidence of low oxygen concentrations occurred not in the surface floodwater plume but in the underlying thermocline layer. Surprisingly, the enrichment of several oxygen minimum zone (OMZ)-associated taxa was not localized near the mortality site but rather occurred widely both throughout the deep portions of the water column and across both the East and West Flower Garden Banks.

## 2. MATERIALS AND METHODS

### 2.1 Site Description

Designated in 1992 as a national marine sanctuary, the Flower Garden Banks are a pair of ocean banks, West Bank and East Bank, located in the northwestern Gulf of Mexico approximately 190 km offshore of Galveston, Texas (Fig. 2), on the edge of the continental shelf. Created by underlying salt domes, each bank rises more than 100 m above the surrounding seabed to within 16 m of the sea surface. The tops of each bank are capped by shallow water coral reef communities that provide critical habitat for ecologically and economically important fisheries in the northwestern Gulf of Mexico.

**Figure 2.**
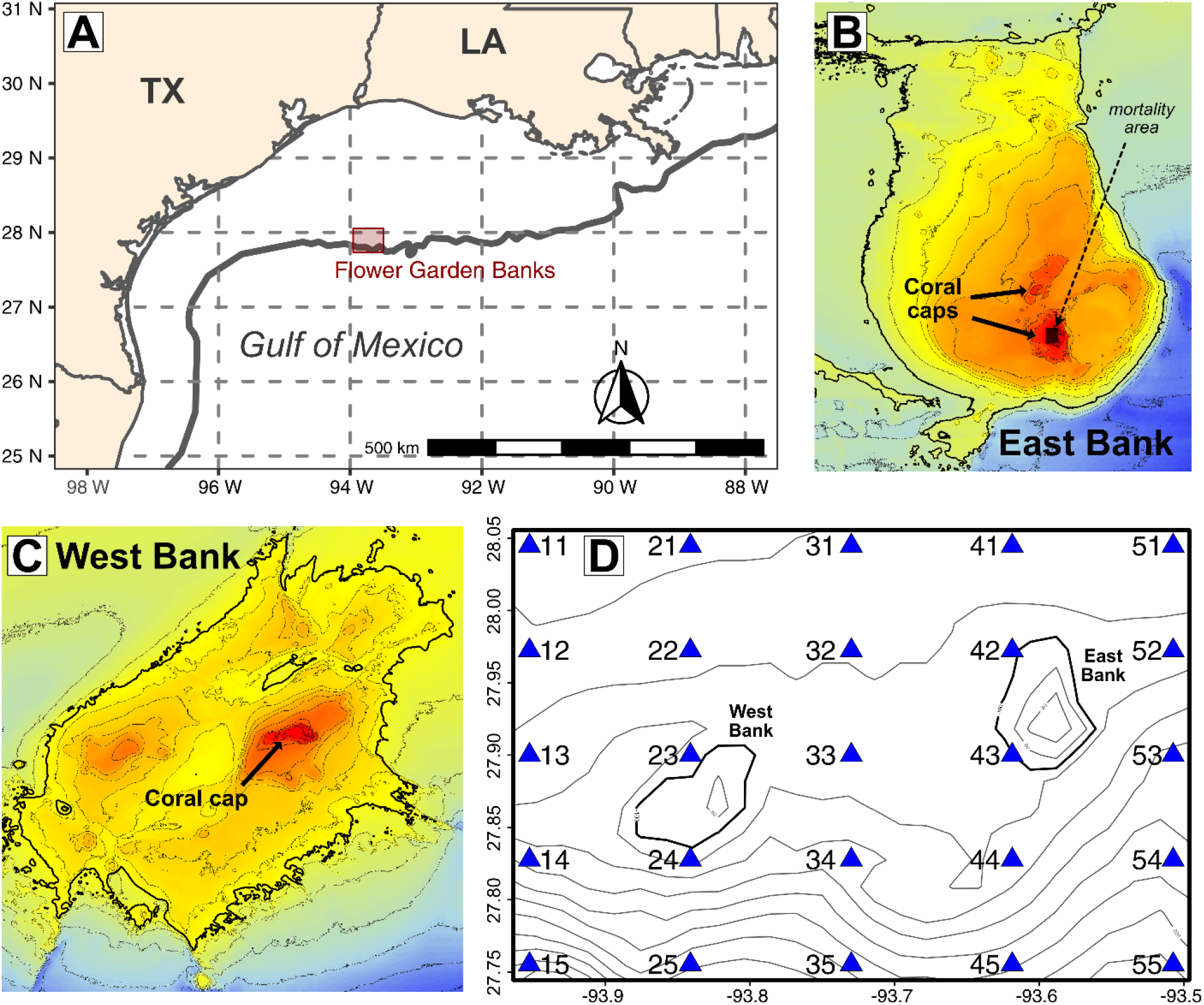
**(A)** Northwestern Gulf of Mexico map with 200-m isobath in gray and location of our sampling grid over the Flower Garden Banks in red. **(B,C)** Multibeam bathymetry data from the USGS are shown over West Bank and East Bank. All lines are 10-m isobath intervals. The 100-m isobath is bolded for reference. The shaded box denotes the location of the 2016 mortality event. **(D)** Map of the sampling grid over the Flower Garden Banks. Location of CTD casts (blue triangles) are labeled by station number. Broad scale bathymetry, shown as gray 20-m isobath intervals, is from the NOAA 1 arc-minute global relief model. The 100-m isobath is bolded for reference.

### 2.2 Sampling

Four cruises were performed to the Flower Garden Banks: 30 July - 02 August 2016 aboard the R/V *Manta*, 20-24 October 2017 aboard the R/V *Point Sur*, 02-07 August 2018 aboard the R/V *Pelican*, and 23-28 October 2018 aboard the R/V *Point Sur*. During each cruise, we collected samples at 25 CTD stations arranged in a 5×5 grid that encompassed the East and West Banks of the Flower Garden Banks (Fig. 2D). The August 2016 cruise occurred as soon as logistically possible (5 days) after the mortality event was discovered on July 25. Seawater samples were collected using a Niskin bottle rosette on a Seabird Electronic (SBE) 25 CTD profiler.

### 2.3 Oxygen and nutrient measurements

Oxygen concentrations were determined *in situ* with a sensor attached to the Niskin rosette and were verified using the Winkler method (12) shipboard. For nutrient measurements, 60mL water samples were filtered through 0.45 μm pore-size filters and frozen at −20°C. Dissolved inorganic nitrate, nitrite, silicate, phosphate, urea, and ammonium were quantified via standard colorimetric analysis using an Astoria autoanalyzer (13).

### 2.4 DNA Extraction and PCR amplification

For each sample, microorganisms were collected from 1 L of seawater by vacuum filtration (≤ 20 cm Hg) onto 47 mm, 0.22 μm Supor PES filter membranes (Pall). Filters were immediately stored at −20°C on the ship after collection. After returning to port, filters were transported on dry ice to Texas A&M University (College Station, TX) and stored at −80°C until DNA extraction. Total DNA was extracted from filters using FastDNA Spin kits (MP Biomedical) and then stored at −20°C until PCR amplification.

The hyper-variable V4 region of the 16S rRNA gene was PCR amplified from DNA extracts using barcoded 515F-806R primer pairs as described previously (3). Amplicon libraries were sequenced at the Georgia Genomics Facility (Athens, GA, USA) using Illumina MiSeq sequencing (v2 chemistry, 2×250bp).

### 2.5 Processing of 16S rRNA amplicon sequences

Raw fastq files were processed using the ‘DADA2’ R package v.1.18 (14). Forward and reverse reads were truncated at lengths of 240 bp and 160 bp, respectively. A maximum expected error threshold of 2 was imposed, and reads were truncated at the first base with a *Q* score of 2 or below before denoising with the DADA2 sample inference algorithm. Denoised reads were merged into amplicon sequence variants (ASVs) using a global ends-free alignment and those containing mismatches in the overlapping region were discarded. ASVs from separate Illumina runs were combined into a single matrix with the *mergeSequenceTables* function. Chimeric ASVs were identified and removed using the consensus method within the *removeBimeraDenovo* function. Taxonomy was assigned using the Silva 138 rRNA database with the IDTAXA algorithm in the ‘decipher’ R package (15). ASVs that were classified as plastids or for which at least a phylum-rank taxonomy could not be assigned were removed from the dataset. Retained ASVs were further screened by aligning them to the Silva 138 rRNA database (16); those which did not properly overlap the targeted V4 hypervariable region or contained a homopolymer >8 bp in length were removed. As a final curation, ASVs that were observed in a sequenced negative control, present in fewer than 2% of total samples, or having a combined relative abundance of less than 0.001% across all samples were removed from analysis.

### 2.6 Ecological analyses

All downstream analyses of the 16S rRNA amplicon data were performed using compositional approaches. Normalization was performed using a robust centered log-ratio transformation followed by matrix completion to impute missing data (17). Dissimilarity between samples was then calculated as the Aitchison distance metric (18). Aitchison distance-based redundancy analysis (db-RDA) was used to summarize the variation in microbial community composition that could be explained by linear relationships with environmental variables. Due to differences in units, all environmental variables were centered and scaled before model calculation. Each db-RDA model was initially constrained by the following variables: individual cruise (categorical), salinity (PSU), temperature (°C), urea, total inorganic nitrogen (TIN), phosphate (HPO_4_^2-^), and dissolved oxygen (mg L^-1^). The significance of the marginal effect of each variable was assessed using PERMANOVA tests (calculated using the ‘vegan’ R package, function ‘anova.cca’) (19). Variables which did not contribute significantly to a model (p>0.05) were dropped and the db-RDA recalculated. Differential abundance analysis was performed using ANCOM-BC (20).

### 2.7 Data Availability

Raw sequence data files are available in the NCBI Sequence Read Archive under accession numbers PRJNA509639 and PRJNA691373.

## 3. RESULTS AND DISCUSSION

### 3.1 Microbial plankton communities varied with water mass

Principal component analysis (PCA) of microbial community beta-diversity (robust Aitchison distances) from all 4 cruises revealed samples were structured by depth (Fig. 3). Given this vertical stratification, we binned samples into two groups: (1) the surface mixed layer, and (2) the underlying thermocline. Comparing these two groups, communities from the mixed layer were significantly less heterogenous (decreased beta dispersion; Fig. 3D) and less diverse (inverse Simpson; Fig. 3C) than those from the thermocline (Mann-Whitney U tests, p<0.01). This pattern is likely a reflection of the comparatively steeper temperature gradients found within the thermocline (ΔT across total depth profile = 8.7 ± 3.6 °C; mean ± SD) versus those found in surface mixed layers (ΔT = 0.7 ± 0.3 °C). Community compositions within the mixed layer were characterized by elevated populations of *Prochlorococcus* (MIT9313) and other *Cyanobacteria*, SAR86, and SAR11 related members (Fig. 4). Combined, these groups accounted for 63.2 ± 13.5% (mean ± SD) of all sequences in the mixed layer. In contrast, communities in the thermocline had larger populations of *Nitrosopelagicus* and other *Thaumarchaeota*, SAR324, and *Thioglobaceae* (*γ-Proteobacteria*) (Fig. 4), which collectively accounted for 29.0 ± 10.7% of all sequences in the thermocline layer.

**Figure 3.**
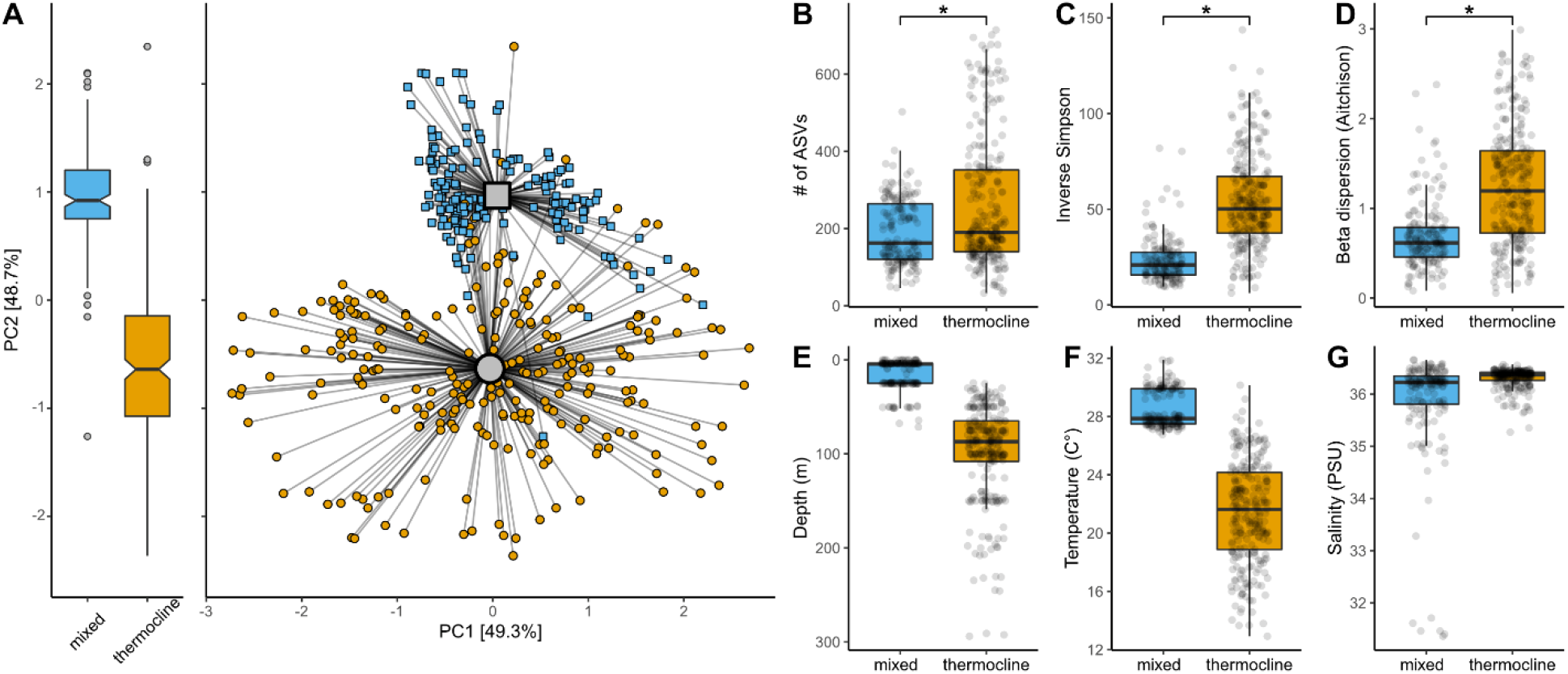
**(A)** A principal-component analysis (PCA) of robust Aitchison distances between 16S rRNA amplicon libraries for all four cruises shows distinct separation between samples collected in the surface mixed layer (blue squares) and those from the underlying thermocline (orange circles). The gray square and circle indicate the respective centroids of mixed layer and thermocline samples. Left boxplots highlight differences between depth layers along the second PC. **(B)** ASV richness was slightly lower in the mixed layer while microbial communities within the thermocline were generally more diverse **(C)** and variable **(D)** in composition (i.e., beta dispersion). The distribution of sample depth, temperature, and salinity are shown in panels **E, F**, and **G**, respectively.

**Figure 4.**
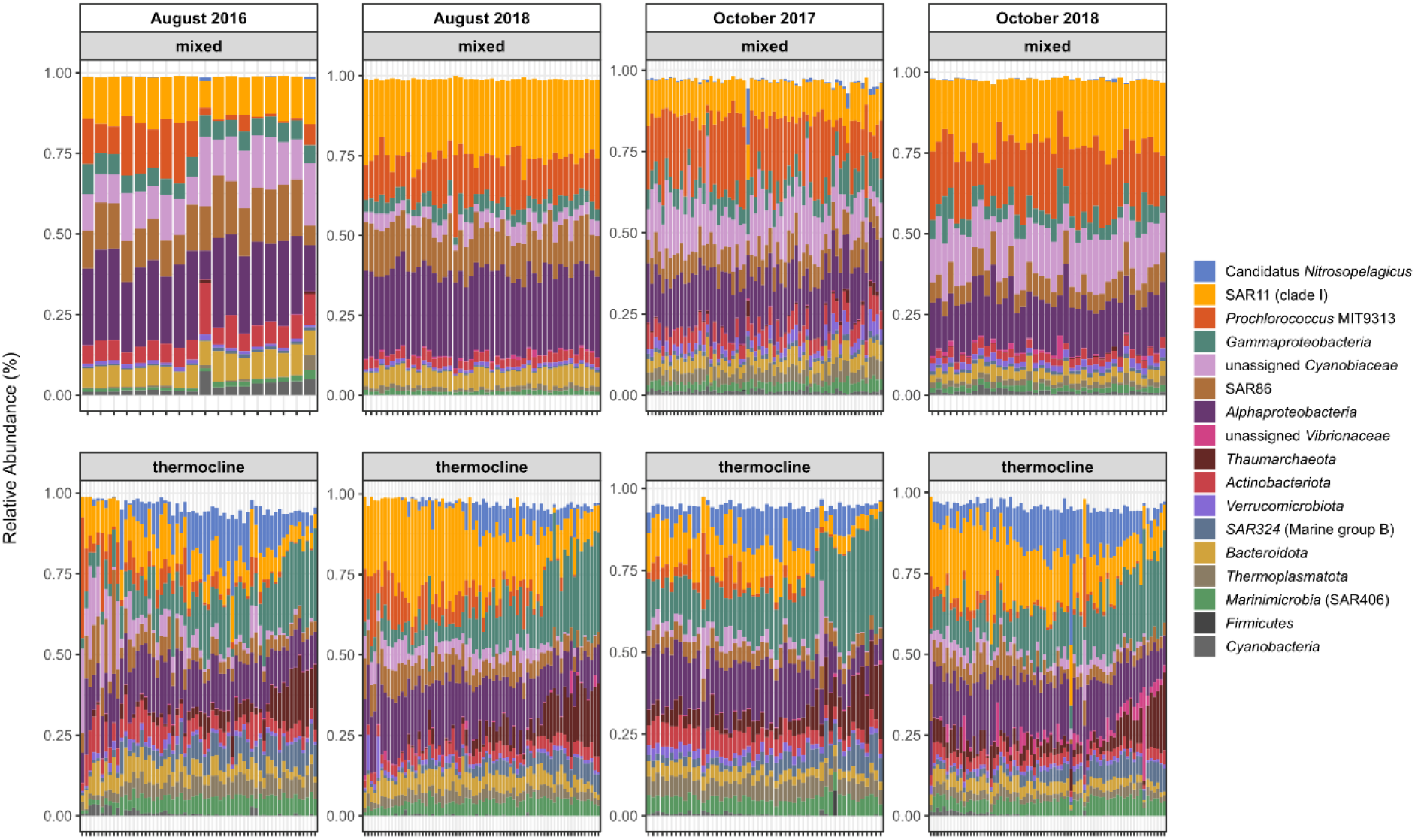
Relative abundances of microbial lineages in seawater samples from the Flower Garden Banks. Within each plot, samples are arranged left to right by increasing depth. Due to the large number of samples, no labels are given on the x-axis for ease of visual interpretation. The plot was constructed to display the highest resolution classification for the most abundant taxa: First, ASVs were clustered at the genus level and any genera having a relative abundance ≥25% in at least one sample were plotted. This procedure was subsequently repeated with the remaining unplotted ASVs at the taxonomic ranks of family, order, class, and phylum. Any remaining rare ASVs left after this procedure were not plotted and contribute to the white space above vertical bars.

### 3.2 Mixed layer microbial communities were strongly affected by a floodwater plume during the mortality event

In the mixed layer, approximately 40% of the compositional heterogeneity among microbial plankton communities was explained by cruise during which they were sampled. Although overall microbial community variability was lower in the mixed layer (Fig. 3D), samples formed distinct clusters for each cruise in a distance-based redundancy analysis (db-RDA; Fig. 5). This pattern indicates there were clear differences in community composition between all four cruises, despite the decreased beta-dispersion. During baseline conditions (i.e. October 2017, August 2018, and October 2018), these differences were generally small and were primarily a result of minor (~4%) variations in the relative abundances of various SAR11 and *Cyanobacteria* ASVs (Table S1). In contrast, the mixed layer microbial communities collected in August 2016—just 5-8 days after the localized mortality event—were notably distinct (Pseudo-F(1,159)=153.6, p<0.001) in RDA space from the other three cruises.

**Figure 5.**
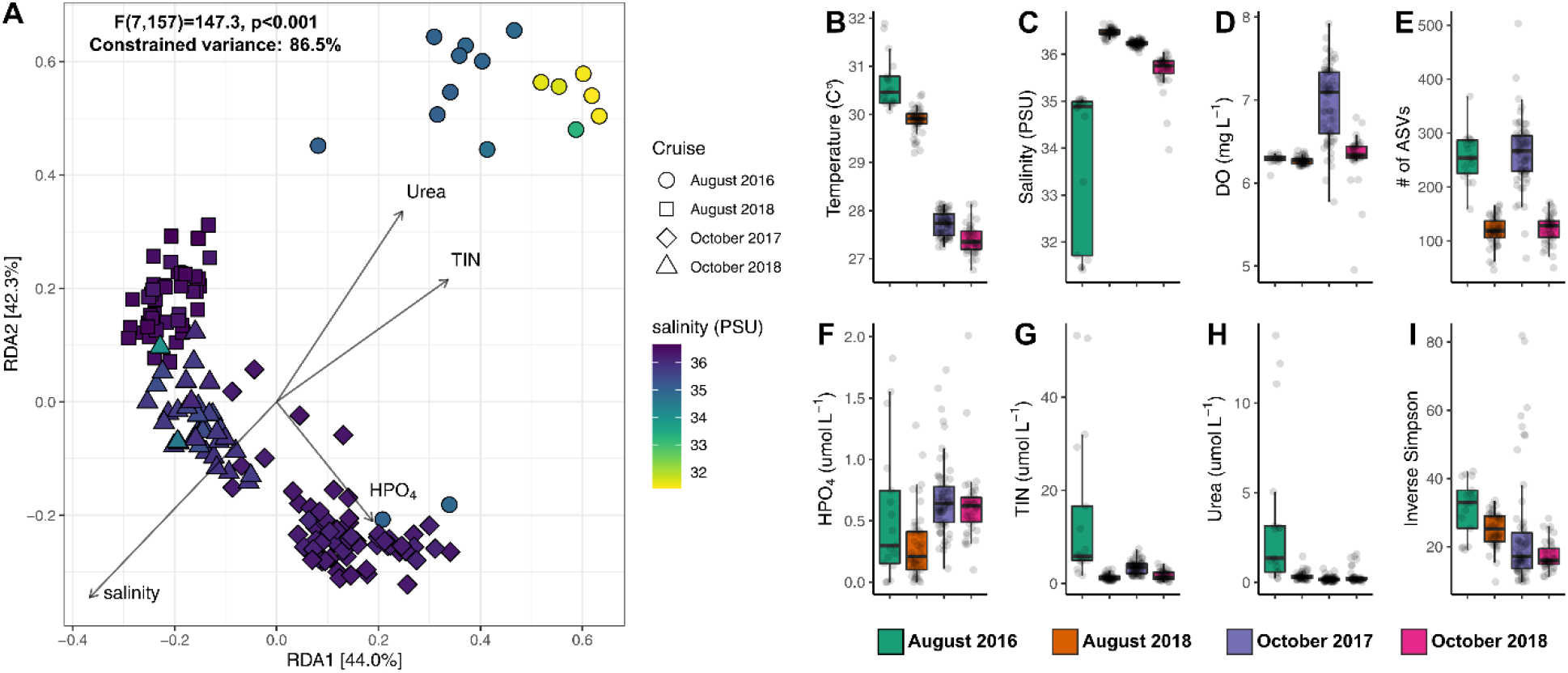
**(A)** Aitchison distance-based redundancy analysis shows microbial communities within the mixed layer during the August 2016 (A16) cruise were distinct from those observed during the other three cruises. These differences coincided with decreased salinity **(C)** and increased concentrations of total inorganic nitrogen **(G)** and urea **(H)**. The remaining panels show the distribution of sample temperature **(B)**, dissolved oxygen concentration **(D)**, ASV richness **(E)**, HPO4 concentration **(F)**, and ASV diversity **(I)**.

In the August 2016 samples, relative abundances of ASVs classified in the SAR11 clade III subgroup were ~18-fold higher than during the other three cruises (Fig. 6). The increase in abundance of this SAR11 subgroup, which typically inhabits brackish or freshwater (21, 22), indicates that microbial community composition and structure within the mixed layer at both West and East Banks were altered by the presence of a floodwater plume shortly after the mortality event. Indeed, satellite imagery from June and July 2016 indicate a surface plume of coastal flood water had reached the Flower Garden Banks (2) prior to the localized mortality event. CTD measurements of near-surface salinities taken during the August 2016 cruise subsequently found this coastal flood water mass was primarily distributed over the East Bank and contained elevated concentrations of total inorganic nitrogen (TIN) and urea (Fig. 5G, H). In agreement with this, we found the highest abundances of SAR11 clade III subgroup in the eastern stations, where salinities were most depressed during the cruise. Notably, stations with the lowest surface salinities and highest abundance of SAR11 clade III subgroup were closest to the area of the East Bank where invertebrate mortality occurred (1). However, this observed spatial west-east pattern was likely only due to timing. Both TABS buoy data (Kealoha et al. 2020) and satellite observations indicate the floodwater plume was transiting the Flower Garden Banks sanctuary from the west and was over the West Bank before our August 2016 cruise first arrived on station (2). Hence, while the floodwater plume was located primarily over the East Bank during the August 2016 cruise, the plume likely impacted mixed layer microbial communities at both banks.

**Figure 6.**
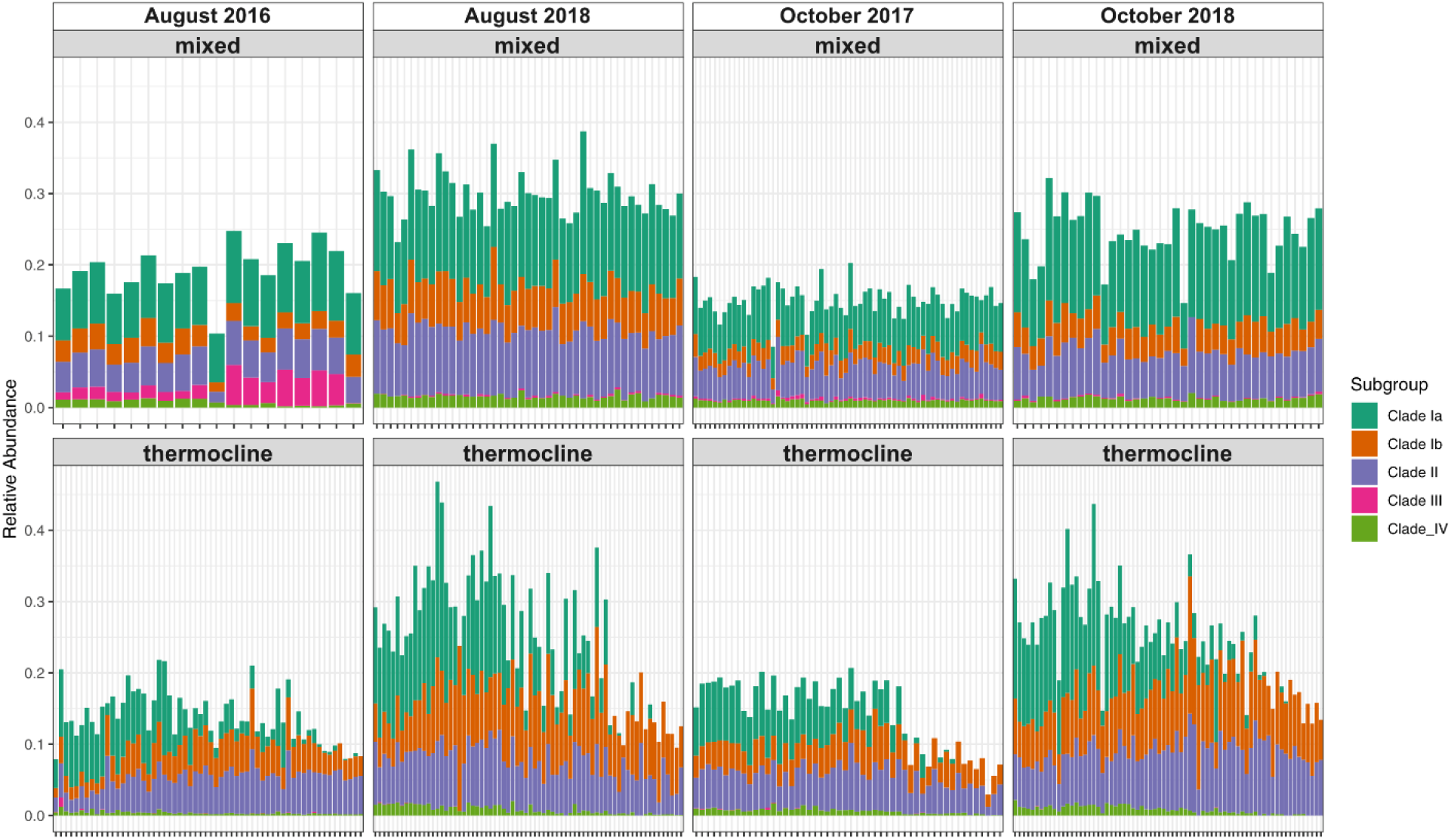
Distribution of SAR11 subgroups in the mixed and thermocline layers across all four cruises. Top panels show data collected from the mixed layer and bottom panels show thermocline data. Within each plot, samples are arranged left to right by increasing depth. Due to the large number of samples, no labels are given on the x-axis for ease of visual interpretation.

### 3.3 Thermocline microbial communities were distinct from baseline conditions during the mortality event

Following the pattern observed in the mixed layer, microbial plankton communities in the underlying thermocline waters during baseline conditions all clustered together in a db-RDA while those collected shortly after the localized mortality event were significantly different (Pseudo-F(1,236)=34.7, p<0.001) (Fig. 7). These differences were not restricted to shallower sections of the thermocline immediately underneath the surface mixed layer—where the floodwater plume was located—but were instead present throughout the entire water column, even in water up to ~300m deep. These differences were also not localized to samples collected near the East Bank, where the mortality event occurred. Instead, we found distinct thermocline communities across the entire 25km × 25km study area. Indeed, unlike in the mixed layer, we did not observe any obvious spatial west-east pattern in the thermocline communities that mirrored the eastern distribution of the floodwater plume during the August 2016 cruise.

**Figure 7.**
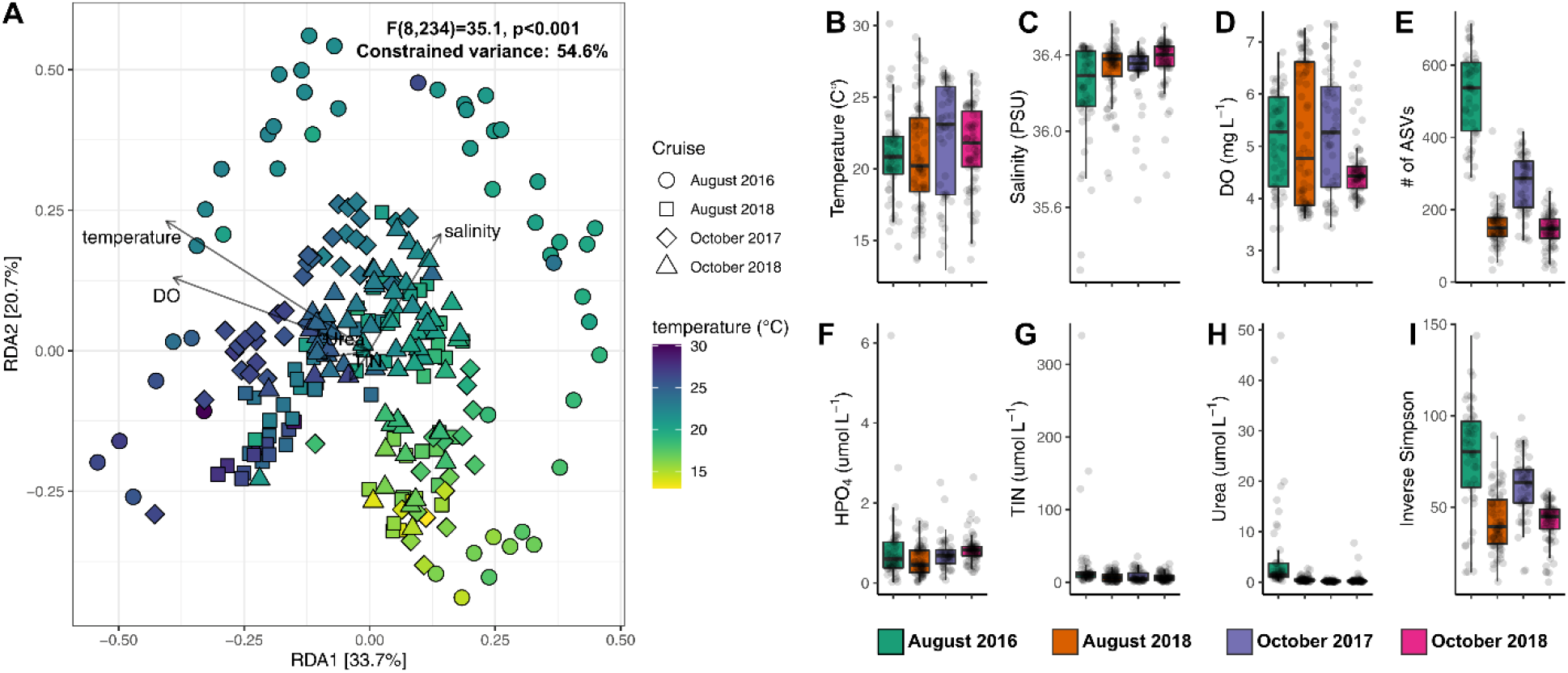
**(A)** Aitchison distance-based redundancy analysis shows microbial communities within the thermocline during the August 2016 cruise were distinct from those observed during the other three sampling periods. These August 2016 thermocline communities had increased ASV richness **(E)** and diversity **(I)** compared to baseline conditions. Despite this, temperature **(B)**, salinity **(C)**, dissolved oxygen concentration **(D)**, HPO4 concentration **(F)**, and concentrations of total inorganic nitrogen **(G)** or urea **(H)** were comparable across all four cruises.

Compared to baseline conditions, changes in thermocline microbial communities 5-8 days after the mortality event appears to have been largely driven by increased ASV richness (Fig. 7E). However, very little of this increased richness was well explained by any measured physiochemical parameters associated with the overlying floodwater plume. Nutrient concentrations within the thermocline were not significantly different from those measured during baseline conditions, and the marginal effects of urea, TIN, and DO concentrations on microbial community structure explained less than ~1% of the microbial community variation we observed across the four cruises. Likewise, although temperature and salinity had larger marginal effects of 8.3% and 11.8%, respectively, neither was significantly different among cruises. Rather, both parameters reflected the typical temperature and salinity gradients found through a thermocline.

### 3.4 Differentially abundant taxa in the thermocline after the 2016 mortality event suggest dissolved oxygen was depressed across both the East and West Banks

Given the weak correlations between microbial community structure and environmental factors in the thermocline samples, we applied a differential abundance analysis to the ASV dataset to determine which taxa were driving the observed differences between the August 2016 cruise and the other three cruises. Bacterial ASVs belonging to the SAR324 clade and *Thioglobaceae* (SUP05 clade) as well as several ASVs of marine ammonia-oxidizing archaea (*Nitrosopumilus* and *Nitrosopelagicus*) were enriched in thermocline samples during the mortality event (Fig. 8). These taxa are commonly found throughout the marine water column but are particularly abundant and active in OMZs, both on continental shelves and in deep water (23–29). We also observed elevated relative abundances of Marine Group II (MGII) *Thermoplasmata*. Although not directly associated with OMZs, phylogenomic analysis of MGII metagenome-assembled genomes suggest this group of Archaea are adapted to oxygen limitation (30). In addition, we observed relative abundances of SAR86 bacteria were depleted within the Flower Garden Banks shortly after the mortality event. SAR86 are an abundant and cosmopolitan marine clade that typically thrive in oxygen-rich (≥70 μM) waters (31, 32). As we did not measure any hypoxia (i.e. DO < 2 mg/L) when these samples were collected, these findings collectively suggest oxygen concentrations were depressed within the thermocline across the entire sampling area some time before the August 2016 cruise. Because that cruise occurred 5-8 days after the 2016 mortality event was first detected, we hypothesize that oxygen concentrations were depressed compared to baseline conditions across both West and East Bank subsurface thermocline waters. Whether these waters were hypoxic is not known.

**Figure 8.**
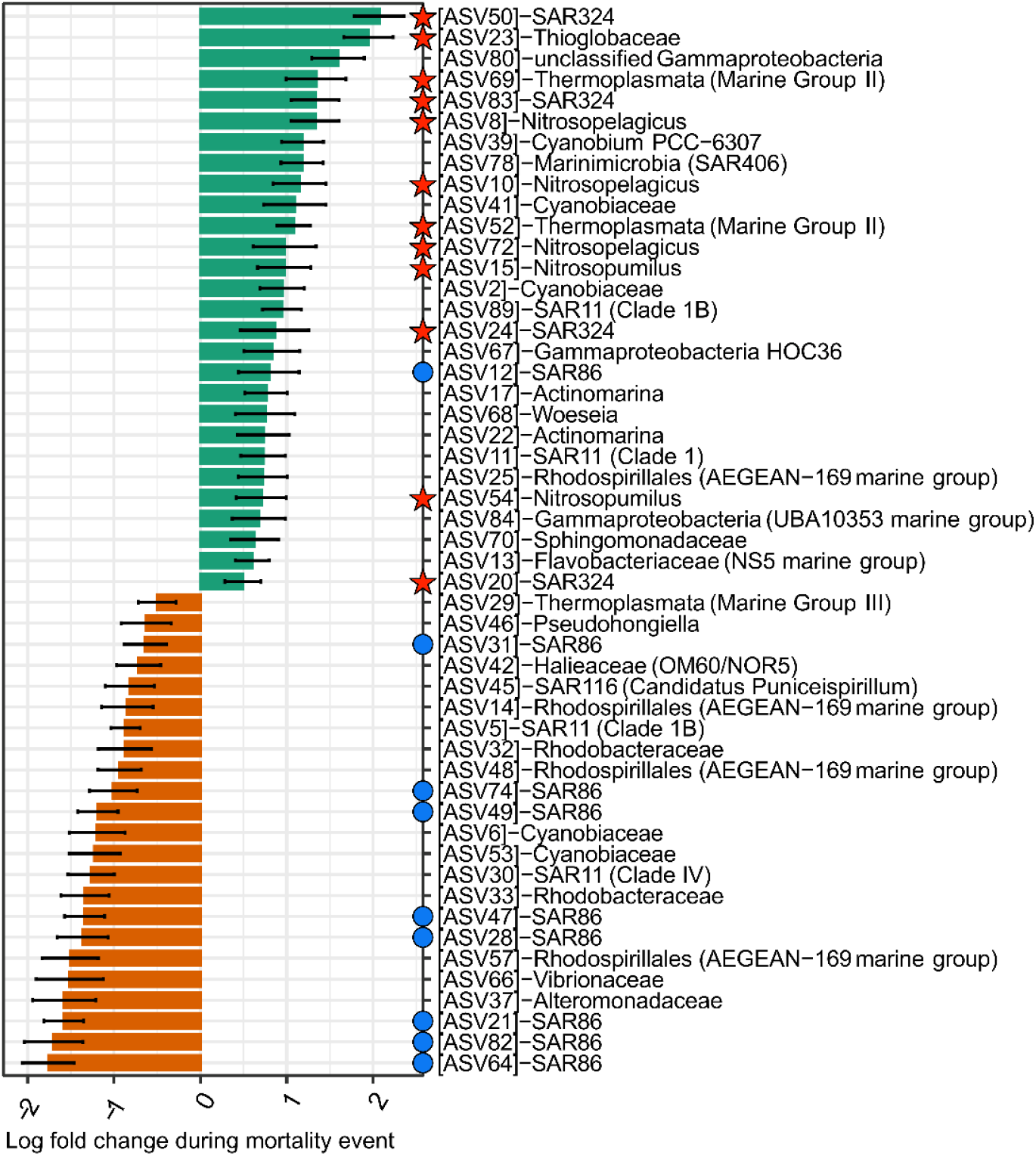
Differentially abundant ASVs within thermocline samples at the Flower Garden Banks. ASVs shown in green were more abundant shortly after the 2016 mortality event. ASVs in orange were more abundant during baseline conditions. Red stars indicate ASVs belonging to taxa which are known to be active and abundant in oxygen minimum zones or have known adaptations to oxygen limitation. Blue circles highlight taxa which are typically found in oxygen-rich waters.

The specific mechanisms that could have lowered oxygen concentrations so far offshore remain unclear. One possible explanation is that an increased flux of organic material sinking from the mixed layer drove increased respiration rates in the thermocline. Although we did not measure organic matter in the water column, a microbiome analysis of both diseased and visually healthy sponges samples collected from the East Bank after the mortality event found the presence of several human fecal indicator bacteria within the sponges’ tissues (33). This shows organic material originating in the overlying floodwater plume had made its way into the thermocline. Respiration of plume-derived organic matter could then plausibly lead to depressed oxygen concentrations across the East and West Bank, similar to the mechanism by which seasonal bottom hypoxia forms on the Texas-Louisiana shelf (2, 3, 34, 35) during the summer. One issue with this hypothesis is that these seasonal low-oxygen waters are typically found inshore of the 60 m isobath and quickly ventilate within ~10km as they move offshore and encounter oxygen-rich water (8, 9, 36). As a result, seasonal plumes of hypoxic water typically do not extend far enough offshore to impact benthic organisms within the East and West Bank coral reefs. One explanation for the 2016 localized mortality event may thus be that the unusually high amount of coastal precipitation along the Gulf coast in the spring of 2016 (37) led to an abnormally large plume of floodwater which was able to stretch farther offshore than in typical years (2). However, this mechanism alone would not explain why the mortality event only occurred in a relatively small area of the East Bank coral reef. Indeed, the lack of any observed mortality at the West Bank reef, which was also impacted by the floodwater plume, strongly suggests an additional mechanism such as the upwelling of low-oxygen deep water was involved.

### 3.5 The East Bank’s coral reef was within the thermocline layer during the mortality event

The depth of the surface mixed layer at each station was calculated from CTD profiles using a temperature difference criterion of 0.5 °C (38)(Fig. S1). Seawater beneath the mixed layer depth was considered to be within the thermocline, which extended to the seafloor at all stations for all cruises (Fig. S1). The mixed layer was deepest in October 2017 with an average depth of ~68m. Mixed layer depth was on average 29m during the other three cruises. These depths are consistent with those previously observed by an array of temperature/conductivity/pressure (TCP) string moorings in the Flower Garden Banks from December 2010 to December 2011 which showed mixed layer depths between 20-40m for spring, summer, and early fall followed by seasonal deepening to 60-80m in the late fall and winter (5).

This analysis also revealed a critical difference between the West and East bank that may explain why the mortality event was localized to the East Bank coral reef. While the West Bank coral reef was within the mixed layer during all four cruises, the mixed layer depth near the East Bank localized mortality area appears to have shoaled slightly during the August 2016 mortality event cruise, possibly due to upwelling of deeper waters up the southern slope of the bank (Fig. 9). As a result, the East Bank coral reef was uniquely within the underlying thermocline layer in August 2016, where microbial data suggests oxygen had been depressed (Fig. 8).

**Figure 9.**
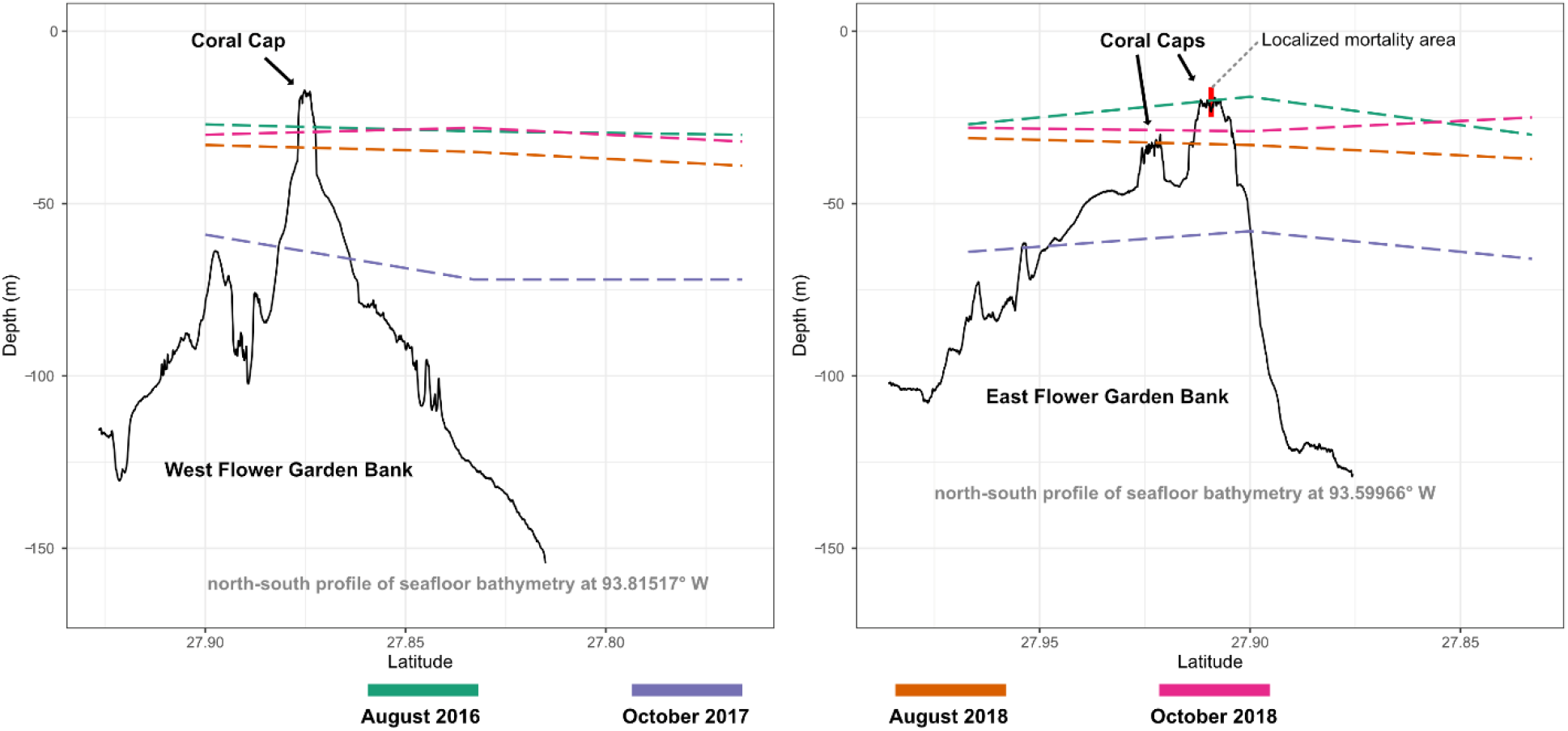
In each panel, the black lines display north-south profiles of the seafloor bathymetry of the West and East Flower Garden Bank coral caps, respectively, based on multibeam bathymetry data from the USGS. Colored dashed lines indicate the bottom of the mixed layer calculated from CTD profiles conducted during each cruise. The red vertical bar in the right-hand panel denotes the localized area of invertebrate mortality that occurred in August 2016. Note, this is the only time when the mortality area was within the underlying thermocline.

### 3.6 Conclusions

The lack of in situ observations of depleted oxygen concentrations previously prevented a definitive conclusion on whether hypoxia was the primary cause of the 2016 localized mortality event at East Bank (1). Collectively, this study reinforces this hypothesis and reveals that this hypoxia was created by a combination of two processes (3). First, leading up to the mortality event, an unusually large discharge of turbid coastal floodwater atypically extended nearly 200km offshore and over the Flower Garden Banks. This plume either deposited additional organic matter to the area or inhibited photosynthesis due to the reduction of sunlight via its turbidity. In either case, this likely resulted in the underlying water column becoming net respiring—even during daylight hours—and may be explanation for the observation of enriched hypoxia-associated microbial taxa in the thermocline across the entire study area. Concurrently with the arrival of the floodwater plume, several lines of evidence suggest deep water had upwelled onto the southern coral cap at East Flower Garden Bank and become trapped in depressions on the reef. This stratification would have prevented re-oxygenation from the surrounding water column and led to localized pockets of hypoxia on the reef.

Our work also highlights the importance of collecting baseline data (39), without which we could have drawn these conclusions. Because microbial plankton communities respond rapidly to physiochemical disturbances such as hypoxia, monitoring of microbial communities on reef systems may be useful indicators of imminent change (40). Indeed, significant changes in microbial communities are known to occur at oxygen concentrations significantly higher than the classic threshold for hypoxia (<2.0 mg O_2_ L^-1^) and thus precede effects on macrofauna (11). Recent work has highlighted mortality events related to hypoxia on coral reefs (41–43), indicating that these events are important stressors in reef ecosystems. Both upwelling and hypoxia are affected by ocean circulation, extreme rainfall events, and decrease in oxygen content, which are all affected by climate change. Understanding how these forces interact in the future will be critical for understanding the future health of the Flower Garden Banks and other coral reefs.

## 4. ACKNOWLEDGEMENTS

This research was supported in part by NSF Award OCE-1800913 to KEFS and JBS, OCE-1800904 to SWD, OCE-1800905 to LSV, and OCE-1800914 to ASC. ASC and SWD were additionally supported by NAS Gulf Research Program Early Career Fellowships. We also thank the NSF-REU summer research program (NSF OCE-1455851) for support of J.H. We thank Marissa Nuttall, John Embesi, and James MacMillan (NOAA), Jake Emmert and Kaitlin Buhler (Moody Gardens), and Ryan Eckert (Florida Atlantic University) for field and logistical support. We would also like express our sincere gratitude to all members of the crew and shore support teams of the R/V Manta, R/V Pelican, and R/V Point Sur as well as members of the NOAA FGBNMS office and Moody Gardens Aquarium (Galveston, TX) for facilitating successful cruises.

## SUPPLEMENTAL MATERIAL LEGENDS

**Figure S1.** Temperature profiles at each station in the study area during each cruise. Horizontal dashed lines denote the calculated mixed layer depth.

**Table S1.** ASV taxonomy, read abundances, and sequences. Cruise and depth (m) are displayed above each sample.

## REFERENCES

1. Johnston MA, Nuttall MF, Eckert RJ, Blakeway RD, Sterne TK, Hickerson EL, Schmahl GP, Lee MT, MacMillan J, Embesi JA. 2019. Localized coral reef mortality event at East Flower Garden Bank, Gulf of Mexico. Bulletin of Marine Science 95:239–250.

2. Le Hénaff M, Muller-Karger FE, Kourafalou VH, Otis D, Johnson KA, McEachron L, Kang H. 2019. Coral mortality event in the Flower Garden Banks of the Gulf of Mexico in July 2016: Local hypoxia due to cross-shelf transport of coastal flood waters? Continental Shelf Research 190:103988.

3. Kealoha AK, Doyle SM, Shamberger KEF, Sylvan JB, Hetland RD, DiMarco SF. 2019. Localized hypoxia may have caused coral reef mortality at the Flower Garden Banks. Coral Reefs https://doi.org/10.1007/s00338-019-01883-9.

4. Breaker BK, Watson KM, Ensminger PA, Storm JB, Rose CE. 2016. Characterization of peak streamflows and flood inundation of selected areas in Louisiana, Texas, Arkansas, and Mississippi from flood of March 2016. US Geological Survey.

5. Teague WJ, Wijesekera HW, Jarosz E, Fribance DB, Lugo-Fernández A, Hallock ZR. 2013. Current and hydrographic conditions at the East Flower Garden Bank in 2011. Continental Shelf Research 63:43–58.

6. Hetland RD, DiMarco SF. 2008. How does the character of oxygen demand control the structure of hypoxia on the Texas–Louisiana continental shelf? Journal of Marine Systems 70:49–62.

7. Zhang W, Hetland RD, Ruiz V, DiMarco SF, Wu H. 2020. Stratification duration and the formation of bottom hypoxia over the Texas-Louisiana shelf. Estuarine, Coastal and Shelf Science 238:106711.

8. Bianchi TS, DiMarco SF, Cowan JH, Hetland RD, Chapman P, Day JW, Allison MA. 2010. The science of hypoxia in the Northern Gulf of Mexico: A review. Science of The Total Environment 408:1471–1484.

9. Zhang W, Hetland RD, DiMarco SF, Fennel K. 2015. Processes controlling mid-water column oxygen minima over the Texas-Louisiana shelf. Journal of Geophysical Research: Oceans 120:2800–2812.

10. Glasl B, Webster NS, Bourne DG. 2017. Microbial indicators as a diagnostic tool for assessing water quality and climate stress in coral reef ecosystems. Mar Biol 164:91.

11. Spietz RL, Williams CM, Rocap G, Horner-Devine MC. 2015. A Dissolved Oxygen Threshold for Shifts in Bacterial Community Structure in a Seasonally Hypoxic Estuary. PLOS ONE 10:e0135731.

12. Winkler LW. 1888. Die bestimmung des im wasser gelösten sauerstoffes. Berichte der deutschen chemischen Gesellschaft 21:2843 –2854.

13. Gordon L, Jennings J, Ross A, Krest J. 1994. World ocean circulation experiment. WOCE operations manual. Volume 3. The observational programme. Section 3.1. WOCE hydrographic programme. Part 3.1. 3. WHP operations and methods.(revision 1). Woods Hole Oceanographic Institution, MA (United States). WOCE Hydrographic.…

14. Callahan BJ, McMurdie PJ, Rosen MJ, Han AW, Johnson AJA, Holmes SP. 2016. DADA2: High-resolution sample inference from Illumina amplicon data. Nature Methods 13:581–583.

15. Murali A, Bhargava A, Wright ES. 2018. IDTAXA: a novel approach for accurate taxonomic classification of microbiome sequences. Microbiome 6:140.

16. Quast C, Pruesse E, Yilmaz P, Gerken J, Schweer T, Yarza P, Peplies J, Glockner FO. 2013. The SILVA ribosomal RNA gene database project: improved data processing and web-based tools. Nucleic Acids Research 41:D590–D596.

17. Martino C, Morton JT, Marotz CA, Thompson LR, Tripathi A, Knight R, Zengler K. A Novel Sparse Compositional Technique Reveals Microbial Perturbations. mSystems 4:e00016–19.

18. Pawlowsky-Glahn V, Egozcue JJ, Tolosana-Delgado R. 2015. Modeling and analysis of compositional data. John Wiley & Sons.

19. Legendre P, Oksanen J, Braak CJF ter. 2011. Testing the significance of canonical axes in redundancy analysis. Methods in Ecology and Evolution 2:269–277.

20. Lin H, Peddada SD. 2020. Analysis of compositions of microbiomes with bias correction. Nat Commun 11:3514.

21. Morris RM, Vergin KL, Cho J-C, Rappé MS, Carlson CA, Giovannoni SJ. 2005. Temporal and spatial response of bacterioplankton lineages to annual convective overturn at the Bermuda Atlantic Time-series Study site. Limnology and Oceanography 50:1687–1696.

22. Brown MV, Lauro FM, DeMaere MZ, Muir L, Wilkins D, Thomas T, Riddle MJ, Fuhrman JA, Andrews-Pfannkoch C, Hoffman JM, others. 2012. Global biogeography of SAR11 marine bacteria. Molecular systems biology 8:595.

23. Wright JJ, Konwar KM, Hallam SJ. 2012. Microbial ecology of expanding oxygen minimum zones. Nat Rev Microbiol 10:381–394.

24. Forth M, Liljebladh B, Stigebrandt A, Hall POJ, Treusch AH. 2015. Effects of ecological engineered oxygenation on the bacterial community structure in an anoxic fjord in western Sweden. ISME J 9:656–669.

25. Walsh DA, Zaikova E, Howes CG, Song YC, Wright JJ, Tringe SG, Tortell PD, Hallam SJ. 2009. Metagenome of a Versatile Chemolithoautotroph from Expanding Oceanic Dead Zones. Science 326:578–582.

26. Bouskill NJ, Eveillard D, Chien D, Jayakumar A, Ward BB. 2012. Environmental factors determining ammonia-oxidizing organism distribution and diversity in marine environments. Environmental Microbiology 14:714–729.

27. Peng X, Jayakumar A, Ward BB. 2013. Community composition of ammonia-oxidizing archaea from surface and anoxic depths of oceanic oxygen minimum zones. Front Microbiol 0.

28. Qin W, Meinhardt KA, Moffett JW, Devol AH, Armbrust EV, Ingalls AE, Stahl DA. 2017. Influence of oxygen availability on the activities of ammonia-oxidizing archaea. Environmental Microbiology Reports 9:250–256.

29. Ulloa O, Canfield DE, DeLong EF, Letelier RM, Stewart FJ. 2012. Microbial oceanography of anoxic oxygen minimum zones. Proceedings of the National Academy of Sciences 109:15996– 16003.

30. Rinke C, Rubino F, Messer LF, Youssef N, Parks DH, Chuvochina M, Brown M, Jeffries T, Tyson GW, Seymour JR, Hugenholtz P. 2019. A phylogenomic and ecological analysis of the globally abundant Marine Group II archaea (Ca. Poseidoniales ord. nov.). ISME J 13:663–675.

31. Aldunate M, De la Iglesia R, Bertagnolli AD, Ulloa O. 2018. Oxygen modulates bacterial community composition in the coastal upwelling waters off central Chile. Deep Sea Research Part II: Topical Studies in Oceanography 156:68–79.

32. Dupont CL, Rusch DB, Yooseph S, Lombardo M-J, Alexander Richter R, Valas R, Novotny M, Yee-Greenbaum J, Selengut JD, Haft DH, Halpern AL, Lasken RS, Nealson K, Friedman R, Craig Venter J. 2012. Genomic insights to SAR86, an abundant and uncultivated marine bacterial lineage. ISME J 6:1186–1199.

33. Shore A, Sims JA, Grimes M, Howe-Kerr LI, Grupstra CGB, Doyle SM, Stadler L, Sylvan JB, Shamberger KEF, Davies SW, Santiago-Vázquez LZ, Correa AMS. 2021. On a Reef Far, Far Away: Anthropogenic Impacts Following Extreme Storms Affect Sponge Health and Bacterial Communities. Front Mar Sci 0.

34. Rabalais NN. 1999. Characterization of Hypoxia: Topic 1, Report for the Integrated Assessment on Hypoxia in the Gulf of Mexico. US Department of Commerce, National Oceanic and Atmospheric Administration.…

35. Turner RE, Rabalais NN. 1994. Coastal eutrophication near the Mississippi river delta. Nature 368:619–621.

36. DiMarco S, Zimmerle H. 2017. MCH atlas: oceanographic observations of the mechanism controlling hypoxia project. Texas Sea Grant TAMU-SG-17-601.

37. Nielsen ER, Schumacher RS. 2020. Dynamical Mechanisms Supporting Extreme Rainfall Accumulations in the Houston “Tax Day” 2016 Flood. Monthly Weather Review 148:83–109.

38. de Boyer Montégut C, Madec G, Fischer AS, Lazar A, Iudicone D. 2004. Mixed layer depth over the global ocean: An examination of profile data and a profile-based climatology. Journal of Geophysical Research: Oceans 109.

39. Martindale RC, Holstein D, Knowlton N, Voss JD, Weiss AM, Correa AM. 2021. Gulf of Mexico Reefs: Past, Present and Future. Gulf of Mexico Reefs: Past, Present and Future 5.

40. Leray M, Wilkins LGE, Apprill A, Bik HM, Clever F, Connolly SR, León MED, Duffy JE, Ezzat L, Gignoux-Wolfsohn S, Herre EA, Kaye JZ, Kline DI, Kueneman JG, McCormick MK, McMillan WO, O’Dea A, Pereira TJ, Petersen JM, Petticord DF, Torchin ME, Thurber RV, Videvall E, Wcislo WT, Yuen B, Eisen JA. 2021. Natural experiments and long-term monitoring are critical to understand and predict marine host-microbe ecology and evolution. PLOS Biology 19:e3001322.

41. Altieri AH, Harrison SB, Seemann J, Collin R, Diaz RJ, Knowlton N. 2017. Tropical dead zones and mass mortalities on coral reefs. PNAS 114:3660–3665.

42. Johnson MD, Scott JJ, Leray M, Lucey N, Bravo LMR, Wied WL, Altieri AH. 2021. Rapid ecosystem-scale consequences of acute deoxygenation on a Caribbean coral reef. Nat Commun 12:4522.

43. Hughes DJ, Alderdice R, Cooney C, Kühl M, Pernice M, Voolstra CR, Suggett DJ. 2020. Coral reef survival under accelerating ocean deoxygenation. Nat Clim Chang 10:296–307.

